# Involvement of c-Jun N-terminus kinase 2 (JNK2) in Endothelin-1 (ET-1) mediated neurodegeneration of retinal ganglion cells

**DOI:** 10.1101/2020.11.03.367151

**Authors:** Bindu Kodati, Dorota L. Stankowska, Vignesh R. Krishnamoorthy, Raghu R. Krishnamoorthy

## Abstract

**Purpose:** The goal of this study was to determine if JNK2 plays a causative role in endothelin-mediated loss of RGCs in mice.

**Methods:** JNK2^-/-^ and wild type (C57BL/6) mice were intravitreally injected in one eye with 1 nmole of ET-1, while the contralateral eye was injected with the vehicle. At two time points (2 h and 24 h) following the intravitreal injections, retinal sections were obtained and phosphorylated c-Jun was assessed. In a separate set of experiments, JNK2^-/-^ and wild type mice were intravitreally injected with either 1 nmole of ET-1 or its vehicle, and euthanized 7 days post-injection. Retinal flat mounts were stained with antibodies to the RGC marker, Brn3a, and surviving RGCs were quantified. Axonal degeneration was assessed by imaging PPD stained optic nerve sections.

**Results:** Intravitreal ET-1 administration produced a significant increase in immunostaining for phospho c-Jun in wild type mice, which was appreciably lower in the JNK2 ^-/-^ mice. A significant (p<0.05) 26% loss of RGCs was found in wild type mice, 7 days post injection with ET-1. JNK2^-/-^ mice showed a significant (p=0.36) protection from RGC loss following ET-1 administration, compared to wild type mice injected with ET-1. A significant decrease in axonal counts and an increase in the collapsed axons was found in ET-1 injected mice eyes.

**Conclusion:** JNK2 appears to play a major role in ET-1 mediated loss of RGCs in mice. Neuroprotective effects of JNK2 following ET-1 administration occur mainly in the soma and not in the axons of RGCs.

## Introduction

The vasoactive endothelin peptide, endothelin-1 (ET-1), has been shown to be elevated in the aqueous humor and circulation in animal models of glaucoma, as well as in glaucoma patients. Several studies have shown the ability of ET-1 administration (intravitreal and retrobulbar) to produce optic nerve degeneration and apoptosis of retinal ganglion cells. However, the detailed cellular and signaling mechanisms contributing to ET-1 mediated neurodegeneration in glaucoma are not completely understood. ET-1 act through two classes of G protein-coupled receptors called ET_A_ and ET_B_ receptors. The regulation of transcription of the preproET-1 gene involves activator protein-1 (AP-1) (complexes of c-Jun and c-Fos), nuclear factor-1 and GATA2 ^[1-3]^. Previously, we demonstrated that c-Jun and its phosphorylation is enhanced in primary RGCs treated with ET-1 ^[4]^. Moreover, ET-1 treatment also produced and increase in ET_A_ and ET_B_ receptor expression in primary RGCs ^[4]^. In addition, c-Jun overexpression was found to produce increase in expression of ET_A_ and ET_B_ receptors in HNPE cells ^[5]^. This suggest that various noxious stimuli including, TNF-α ^[6]^, and hypoxia ^[7]^, that activate c-Jun could trigger the expression of both ET-1 and its receptors. c-Jun is activated by phosphorylation by the stress activated protein kinase, c-Jun N-terminal kinase (JNK) which is activated in response to numerous stimuli relevant to glaucoma pathogenesis.

The stress induced MAPK, c-Jun N-terminal kinase (JNK) is involved in the various cellular activities from proliferation to apoptosis which is mainly dependent on the response of the specific cell type, external stimuli and its activation ^[8]^. There are three isoforms of JNK, JNK1, JNK2 which are ubiquitously expressed in most cells and JNK3 which is expressed mainly in brain, heart and testis ^[9-11]^. JNK regulates the activity by phosphorylation of various downstream factors including the c-Jun which is a component of transcription factor AP-1, ATF2, ATF3. In addition, JNK regulates the activity/expression of other transcription factors including, Elk1, p53, HSF-1 and c-Myc and the members of Bcl2 family, involved in both extrinsic and intrinsic mechanisms of apoptosis ^[12-15]^. Studies on cerebellar granule neurons have shown that the phosphorylation of c-Jun in the nucleus is more critical for the apoptosis than the phosphorylation of JNK in the cytosol ^[16]^. Studies have demonstrated apoptotic effects of JNK isoforms and their reversal by blocking JNK with various agents ^[12, 17-21]^. Our previous studies also showed the involvement of JNK in ET-1 mediated cell death in the primary RGC ^[4]^, but the exact JNK isoforms contributing to ET-1 mediated cell death has not been determined. The present study aimed to elucidate the role of JNK2 in endothelin mediated cell death of RGCs in mice.

## Methods

### Animals

All animal procedures were in accordance with the Association for Research in Vision and Ophthalmology (ARVO) resolution on the use of animals in ophthalmic and vision research and approved by the University of North Texas Health Science Center (UNTHSC) Institutional Animal Care and Use Committee (IACUC) (animal protocol #IACUC-2017-0024). Male and female, Wild type (C57BL/6) and JNK2^-/-^ mice were purchased from Jackson Labs and were matched for age and gender. The animals were housed in rooms where the temperature, humidity and light were controlled. Food and water were available ad libitum.

### Intravitreal Injections of ET-1 or vehicle

Mice were anesthetized by injecting intraperitoneally using an anesthetic cocktail having Xylazine (5.5 mg/kg)/ Ketamine (55 mg/kg)/ Acepromazine (1.1 mg/kg). One eye was injected with the endothelin-1 (ET-1) and contralateral eye was injected with the vehicle using a Hamilton syringe. ET-1 obtained from Bachem (Torrance, CA, USA) was resuspended in the vehicle solution (0.25% glacial acetic acid neutralized to pH 7.0 with NaOH and adjusted to a final concentration of 500µM). A single drop of 0.5% proparacaine hydrochloride (Alcon Laboratories, Inc., Fort Worth, TX, USA) and 1% tropicamide were applied to both eyes. An ultrafine 33.5 gauge disposable needle connected to a 10 microliter Hamilton syringe was be used to intravitreally inject 2 µl of either 500 μM ET-1 or vehicle into vitreous chamber by injecting through the sclera 1mm behind the limbus region. The injection was made slowly, making sure to avoid the lens during insertion of the needle. After the injection, the needle was held in place for 1 minute and gradually withdrawn from the vitreal chamber. Following the injection, triple antibiotic was applied at the injection site and the animals were allowed to recover from the anesthesia. After various time points following the intravitreal injection including, 2 h, 24 h and 7days mice were humanely euthanized by an overdose of 120 mg/kg body weight of Fatal-Plus (pentobarbital).

### Immunohistostaining

Mice were euthanized after the 2 h and 24 h time points, their eyes were enucleated and immediately fixed in 4% paraformaldehyde (PFA) and placed on a shaker at room temperature for 3 hours. After the incubation the fixed retinas were washed with PBS and placed in 70% ethanol and thereafter embedded in paraffin and sectioned. The sections were de-paraffinized using the xylene, and rehydrated with graded descending series (100%, 95%, 90%, 80% and 50% ethanol) and finally with the 1X PBS. Permeabilization with 0.1% sodium citrate and 0.1% Triton-X 100 was carried out for 8 minutes in order to facilitate the subsequent entry of antibodies. To prevent non specific binding of the secondary antibody, the sections were incubated with blocking buffer (5% Normal donkey serum/5% BSA in PBS) for approximately 2-3 h. Following blocking, the sections were incubated in the respective primary antibodies overnight at 4°C. The primary antibodies used for the experiment are: mouse anti phospho-c-Jun (1:50, SC-822, SantaCruz Biotechnology), and goat anti-Brn3a (1:200, SC-31984, SantaCruz Biotechnology). Following primary antibody incubations, the sections were washed three times with 1x PBS and incubated with appropriate secondary antibodies for approximately 1-2 hours at room temperature. Secondary antibodies used in the experiment are donkey anti-mouse Alexa 546 (1:1000 dilution, A10036, Life Technologies, Carlsbad, CA), donkey anti-goat Alexa 488 (1:1000 dilution, A11055, Invitrogen). To assess the non-specific binding of secondary antibodies, Blank sections (in which the primary antibody incubation was omitted) were also carried out followed by secondary antibody incubations.

### Retinal Flat Mount Immunostaining

Seven days following intravitreal injections, the animals were euthanized and the eyes were enucleated and immediately fixed overnight in 4% paraformaldehyde (PFA) at 4°C. After three washes with 1x PBS, retinas were carefully separated from the globe and incubated overnight in blocking buffer (5% normal donkey serum/ 5% BSA in PBS) at 4°C, cuts were made in the four quadrants (superior, inferior, nasal and temporal) and retinal flat mounts were prepared. The retinal flat mounts were then incubated with the primary antibody, goat anti-Brn3a (1:200, SC-31984, SantaCruz Biotechnology) for three days at 4°C. After three washes with 1x PBS, the flat mounts were incubated overnight in the corresponding secondary antibodies: Alexa 488 conjugated donkey anti-goat antibody (1:1000 dilution, A11055, Invitrogen,) at 4°C. After washes, the flat mounts were mounted using Prolong Gold anti-fade (Life Technologies) and images were taken using the Keyence fluorescence microscope (Itasca, IL, USA).

### Quantification of retinal ganglion cell survival

Images of flat mounts were captured using a 20x objective in a Keyence fluorescence microscope. Images were taken at two different eccentricities, located at one-third and two-third of the distance between the optic nerve head and the periphery of the retina. Two images were captured at each eccentricity, in each of the four quadrants, including, the superior, inferior, nasal, and temporal quadrants, for a total of 16 images per retina. RGC counts were determined by a semi-automatic cell counting procedure on ImageJ (Rasband, 1997-2018). Briefly, all the images were first converted to an 8-bit images and run through FTT band pass filter. The images were then run through auto-threshold and converted to binary images. After removing outliers, binary functions like Fill holes and Watershed were applied. Cells were then counted by applying appropriate particle parameters i.e. size (0.01-infinity) and circularity (0.0 to 1). The counting was performed by a masked observer who was unaware of the genotypes and treatment groups of the animals.

### Paraphenylenediamine (PPD) staining of optic nerve for the assessment of axonal damage

Degeneration of axons were examined using PPD staining which stains the myelin around the axons. Briefly, the mice were injected with vehicle and ET-1 (wild type and JNK2^-/-^) and maintained for a week. The mice were then sacrificed; the eyes were enucleated the optic nerves were excised posterior to the globe. The optic nerves were then immediately fixed in 2% paraformaldehyde, 2.5% glutaraldehyde in 0.1 M sodium cacodylate buffer. Before dehydration the optic nerves were transferred to 2% osmium tetroxide in PBS for 1 hour and embedded in Epon. Then cross sections were obtained using the ultramicrotome and stained with 1% PPD. Stained PPD Images were taken in Zeiss LSM 510 META confocal microscope under 100X oil immersion objective. Images were taken covering central and each quadrant of the peripheral optic nerve. The counts were obtained using an automated program in ImageJ software and the total axonal counts are the sum of the axonal counts from all the images. Based on the statistical analysis the neuroprotective / neurodegenerative effects were analyzed further.

### Statistical analysis

The statistical analysis was performed using the GraphPad Prism 7 (GraphPad Software, La Jolla, CA). To compare the data between two groups, an unpaired t-test and for Multiple groups, one-way analysis of variance was used. Statistical significance was determined at P<0.05.

## Results

### Significant upregulation of phospho c-Jun in C57BL/6 wild type compared to JNK2^-/-^ mice

ET-1 treatment has been shown to elevate JNK and it substrate: immediate early gene c-Jun in both primary RGC and the HNPE cells ^[4, 22]^. He et al., (2015) also demonstrated a dramatic increase in immunostaining for phospho-c-Jun following treatment of primary RGCs with 100 µM ET-1 for 24 h. In the present study, the involvement of JNK2 and its effects on ET-1 mediated RGC cell death was assessed *in vivo*. Briefly, wild type mice (C57BL/6) and JNK2^-/-^ mice were injected intravitreally with 500 µM ET-1 in one eye and the vehicle in the contralateral eye. At two different time points (2 h and 24 h) post-injection, the animals were sacrificed and retinal sections were obtained. In order to assess the involvement of JNK2 signaling mechanism following intravitreal ET-1 administration the immunohistochemical analysis of phospho c-Jun was performed in the retinal sections.

At the 2 h time point following ET-1 injection, the wild type mice showed a significant increase in immunostaining for phospho-c-Jun mainly in ganglion cell layer (p=0.046, n=3, **Figure: 1B**), compared to the vehicle treated eyes. There was an appreciable increase (not statistically significant) in immunostaining for phospho-c-Jun in the nerve fiber layer, inner plexiform layer and the inner nuclear layer in the ET-1 treated eyes, compared to vehicle treated eyes in wild type mice (**Figure: 1A**). In the JNK2^-/-^ mice in all the retinal layers there was no appreciable increase in the phospho-c-Jun in the ET-1 treated eyes, compared to the vehicle treated eyes. The integrated densities from confocal z-stack images were analyzed for the immunostaining for phospho-c-Jun and compared between the ET-1 and vehicle treated eyes. There was a significant increase in phospho-c-Jun staining in ET-1 injected wild type eyes compared to JNK2^-/-^ mice (n=3 wild type and n=4 JNK2^-/-^) eyes in the GCL (p=0.0007, **Figure: 1B**), IPL (p=0.0272, **Figure: 1C**) and INL (p=0.0014, **Figure: 1D**).

**Figure 1:**
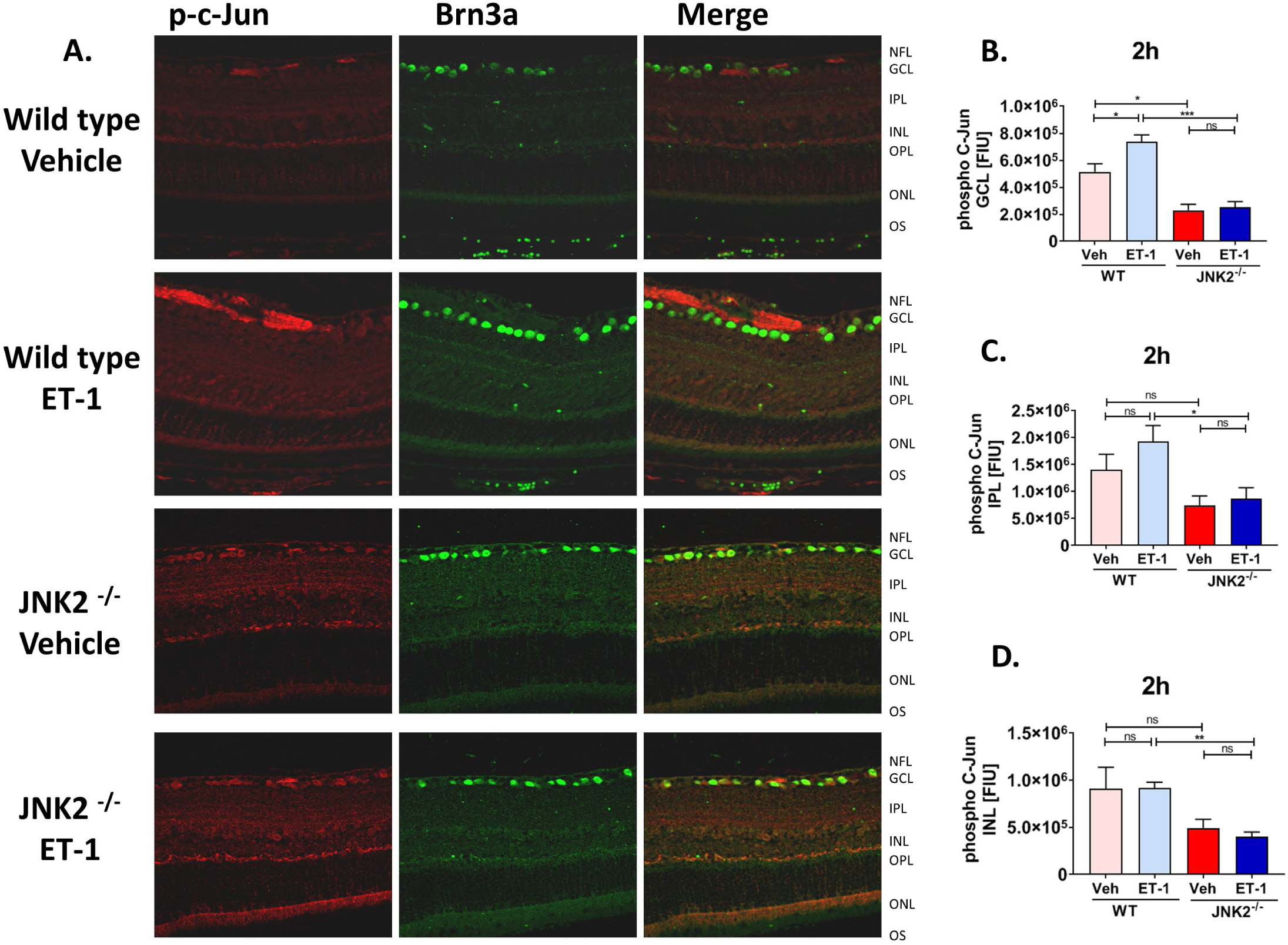
Phospho c-Jun expression in retinas of wild type and JNK2^-/-^ mice 2 h following intravitreal administration of ET-1. A) Representative images from confocal microscopy. Immunostaining of retina sections from wild type and JNK2^-/-^ mice 2 h following intravitreal ET-1 administration with 2µL of 500 µM ET-1 using antibodies against phospho c-Jun (red) and Brn3a (green). Relative fluorescent intensity quantification for GCL (B), IPL (C), and NFL (D) layers respectively. Bars represent mean ± SEM (n = 3 animals in wild type and n=4 in JNK2^-/-^). ONL outer nuclear layer, OPL outer plexiform layer, INL inner nuclear layer, IPL inner plexiform layer, GCL ganglion cell layer, NFL nerve fiber layer. Asterisks indicate statistical significance *p < 0.05; **p < 0.01; ***p< 0.001 unpaired t-test.

At the 24 h time point, wild type mice showed a significant increase in immunostaining for phospho-c-Jun in the GCL (p= 0.0246, n=4, **Figure: 2B**) and INL (p= 0.0247, n=4, **Figure: 2D**) in the ET-1 injected eyes compared to vehicle treated eyes. There was no appreciable increase in immunostaining for phospho-c-Jun in the NFL and IPL in the ET-1 injected eyes, compared to the vehicle injected eyes in wild type mice. In the JNK2^-/-^ mice, there was no significant change in the phospho-c-Jun in all the retinal layers in the ET-1 treated eyes, compared to the vehicle treated eyes. Upon comparison of the ET-1 treated eyes between the wild type and the JNK2^-/-^ mice, we found a significant increase in immunostaining for phospho-c-Jun in the GCL (p=0.0006, n=4, **Figure: 2B**), IPL (p= 0.0022, n=4, **Figure: 2C**) and INL (p=0.0007, n=4, **Figure: 2D**) in the wild type mice, compared to the JNK2^-/-^ mice (**Figure 2A**).

**Figure 2:**
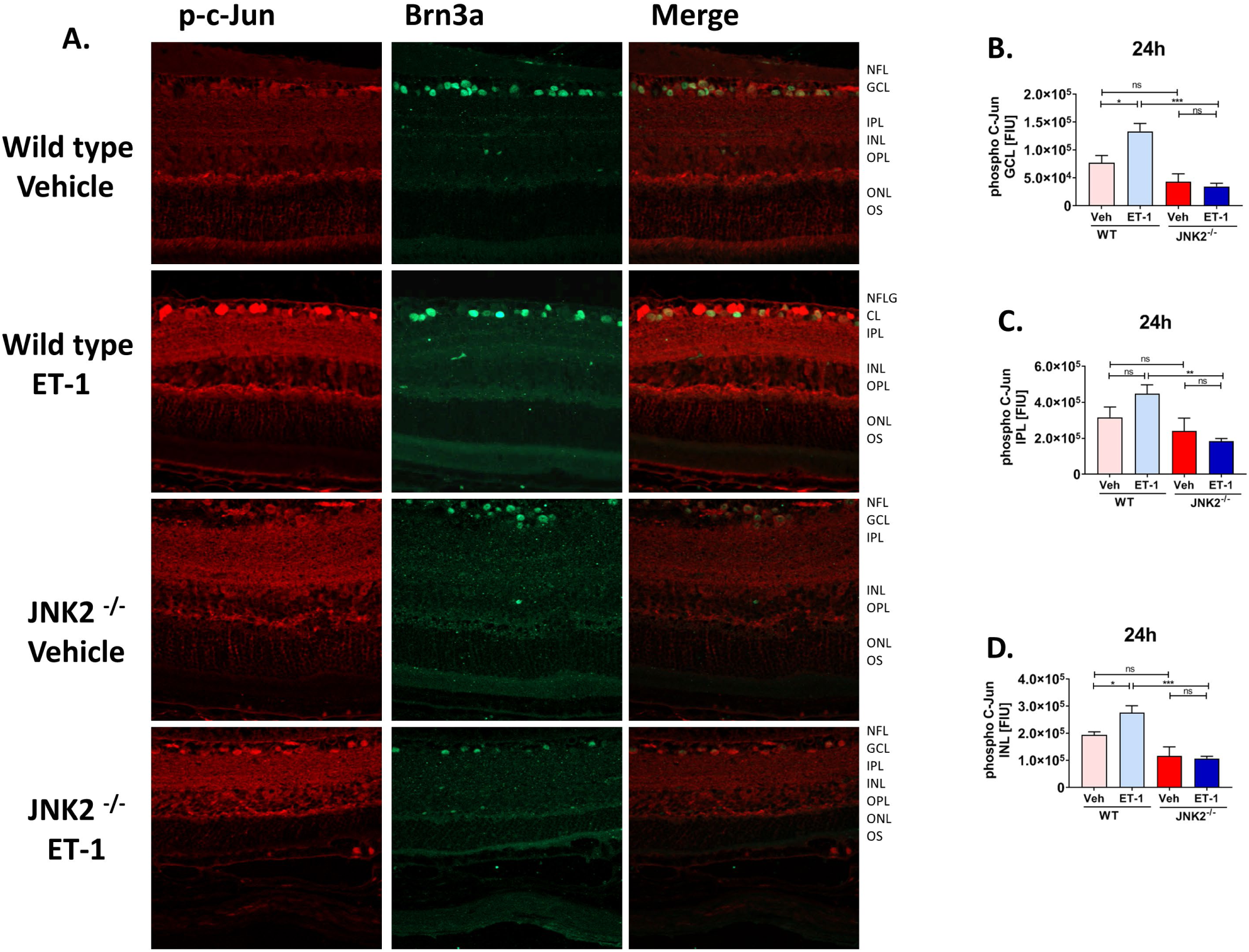
Phospho c-Jun expression in retinas of wild type and JNK2-/-mice 24 h following intravitreal administration of ET-1. A) Representative images from confocal microscopy. Immunostaining of retina sections from mice following 24 h of intravitreal injection with 2 µL of 500 µM ET-1, using antibodies to phospho c-Jun (red) and Brn3a (green). Relative fluorescent intensity quantification for GCL (B), IPL (C), and NFL (D) respectively. Bars represent mean ± SEM (n = 4 animals/group). ONL outer nuclear layer, OPL outer plexiform layer, INL inner nuclear layer, IPL inner plexiform layer, GCL ganglion cell layer, NFL nerve fiber layer. Asterisks indicate statistical significance *p < 0.05; **p < 0.01; ***p< 0.001 unpaired t-test.

### Protection of RGC from apoptosis in JNK2^-/-^ mice in comparison with the wild type mice

To determine the involvement of JNK2 signaling pathway in the loss of RGCs, we intravitreally administered ET-1 in one eye and the vehicle in contralateral eye in both wild type and JNK2^-/-^ mice and maintained for 7 days. Retinal flat mounts were prepared and the survival of RGC were assessed by counting the viable RGC immunolabelled with the Brn3a antibody. The RGC counts between the ET-1 and vehicle administered eyes for each retinal eccentricity was compared between the wild type and JNK2^-/-^ mice. In wild type mice, in the peripheral eccentricity, there was a significant decrease in RGC counts in the ET-1 injected eyes, compared to vehicle injected eyes (**Figure: 3B**). However, there was no statistical significance in RGC counts in the mid-peripheral region between the vehicle-injected and ET-1 injected eyes in wild type mice (**Figure: 3C**). We also found that there is no significant RGC loss in both eccentricities in ET-1 injected eyes, compared to the vehicle injected eyes in JNK2^-/-^ eyes. We observed a statistical significant difference in total RGC counts (peripheral and mid-peripheral eccentricities) between the wild type mice and JNK2^-/-^ mice intravitreally injected with ET-1(**Figure: 3D**). Taken together, the results demonstrate that the JNK2 isoform plays a critical role in signaling in RGC apoptotic death with the ET-1 treatment (**Figure 3A**).

**Figure 3:**
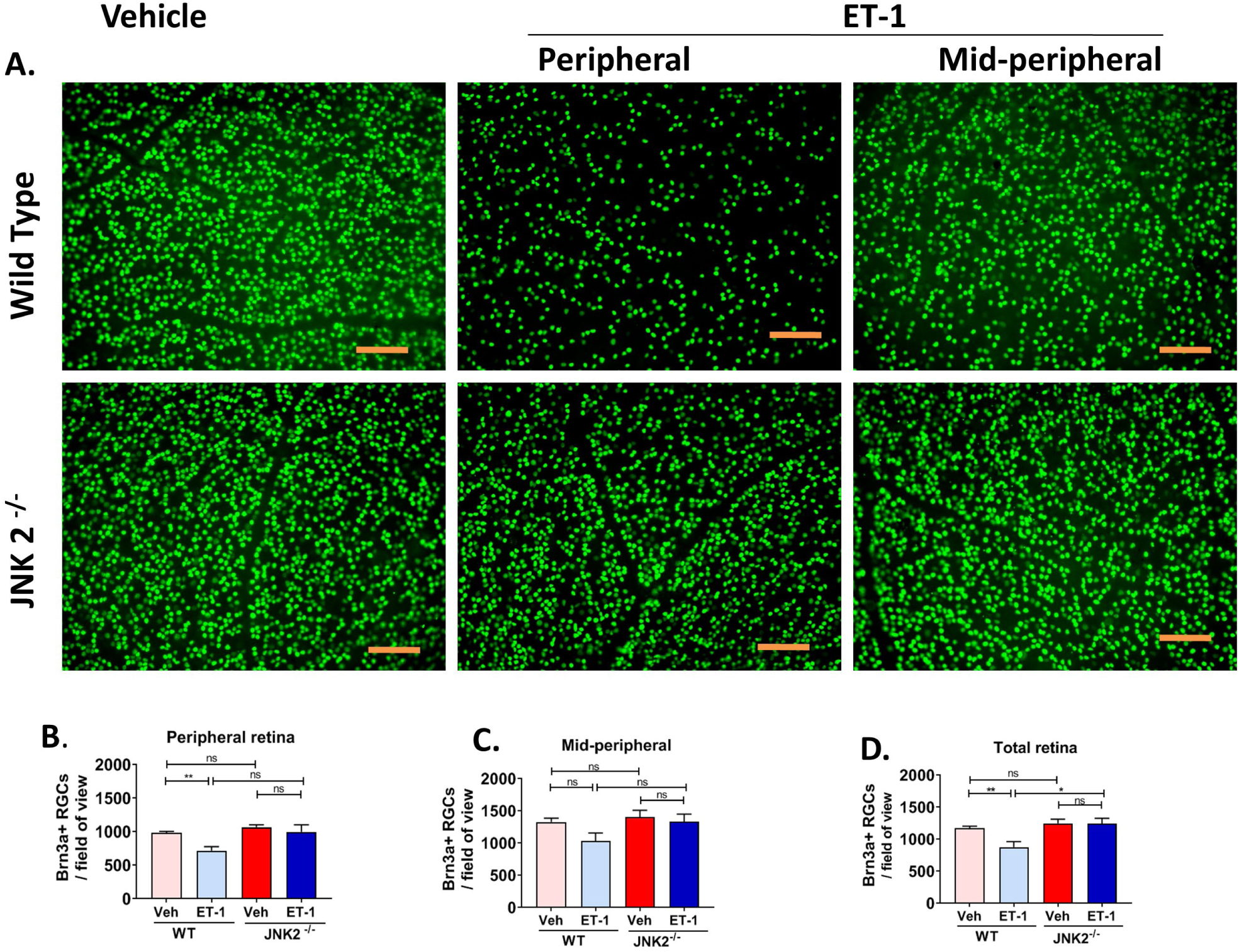
Survival of RGC in JNK2^-/-^ mice compared to C57BL/6 mice following intravitreal administration of ET-1: Mice were Intravitreally injected either with Vehicle or ET-1 and maintained for 7 days. The mice were euthanized and RGC loss was analyzed by the counts obtained from the flat-mounts in two eccentricities (peripheral and mid-peripheral). The vehicle and ET-1 injected images stained with Brn3a marker were represented for the wild-type and JNK2^-/-^ mice for the peripheral and mid-peripheral eccentricities (A). Graphs are shown for peripheral (B) mid-peripheral (C) and total counts (D). Significant loss of RGCs (26% loss) is observed in wild type mice. Data are shown as mean ± SD (n = 6 in wild type and n =5 in JNK2^-/-^, unpaired t-test, two tailed, **p = 0.0082, *p = 0.0156). Scale bar indicated 100µm.

### Axonal degeneration

One week following injection with either ET-1 or vehicle, wild type and the JNK2^-/-^ mice were sacrificed and enucleated. The axonal integrity of the optic nerve axons was assessed by paraphenylenediamine (PPD) staining which darkly stains the myelin of damaged axons. In wild type mice, the optic nerve of ET-1 injected eye showed intense staining of myelin, glial scar formation and disruption of axonal bundles. Wild type mice showed significant disruption of the axonal integrity (based upon the axon counts obtained using Image J) in the ET-1 injected eye (p=0.0154, t test n=7) (**Figure: 4B**), compared the vehicle treated eye. However, when ET-1 injected eyes were compared between the wild type and JNK2^-/-^ mice, there was no significant difference among them. Further we manually counted the degenerative axons which stained darkly in the axoplasm of PPD stained optic nerve cross-sections and analysis of wild type and the JNK2^-/-^ did not show any significant difference. We observed a significant decline in axon counts and increase in collapsed axons in the ET-1 injected eyes, compared to the vehicle injected eyes in wild type (p= 0.0032, n=7) (**Figure: 4B and 4C**). However, there was no significant difference in axon counts and collapsed axons, between wild type and JNK2^-/-^ mice following ET-1 injection (n=5) (**Figure 4**), suggesting that blocking JNK2 signaling does not protect axons from ET-1-mediated degeneration.

**Figure 4:**
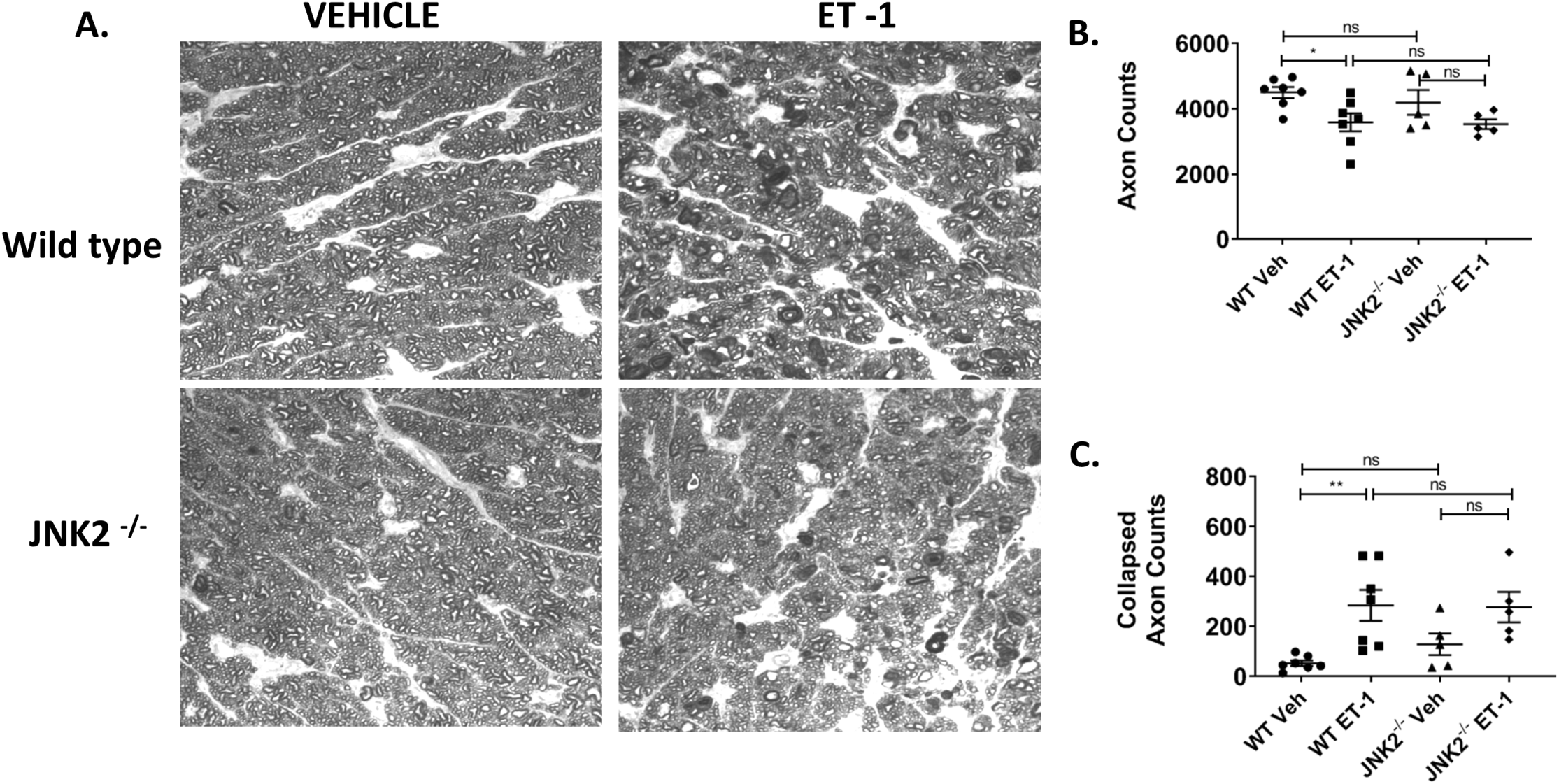
Optic nerve axonal degeneration in wild type and JNK2-/-mice. Both wild type and JNK2-/-mice were intravitreally injected in one eye with vehicle and contralateral eye with ET-1 and maintained for a week. Post injections and maintenance animals were sacrificed and subjected to PPD staining to assess the optic nerve degeneration. A) Severe axonal degeneration along with gliosis and glial scar were observed in ET-1 injected eyes compared to vehicle injected eye in both wild type and JNK2-/-mice. Dark spots indicate the degenerating axons. B) The mean axon counts in ET-1 injected wild type mice showed a significant reduction in number of healthy axons compared to Vehicle injected eyes (*p=0.0154, t test n=7 wild type)). C) The collapsed axons show a significant increase in counts between the ET-1 and vehicle injected eyes in wild type mice. Mean ± SD (n = 7 in Wild type and n =5 in JNK2-/-mice, unpaired t-test, two-tailed, **p = 0.0032). Scale bar indicated 20µm.

## Discussion

The currently available treatments, including, pharmacological or surgical approaches of lowering IOP greatly slow down the progression of the disease ^[23]^. However, in some patients, there is continued neurodegenerative effects. One of the endogenous vasoconstrictors that produces ischemic insult in the ONH, with a potential role in glaucoma and other neurodegenerative diseases is endothelin-1 (ET-1) ^[24, 25]^. *In vivo* studies in rabbits and rhesus monkeys showed that administration of low doses of ET-1 generated ischemic effect leading to the optic nerve damage as in glaucoma ^[26, 27]^. Besides the primary insult of elevated IOP, other factors including, ischemia ^[28]^, excitotoxicity ^[29]^, TNF-α ^[30]^ or a combination of these are thought to be contributors to glaucomatous neurodegeneration. Previous studies have reported the upregulation of ET-1 and its receptors may activate the optic nerve astrocytes, leading to ECM changes in the ONH causing disruption in axoplasmic transport and eventual RGC loss via withdrawal of trophic support and reactive gliosis ^[25, 31-33]^. However, the signaling events contributing to ET-1 mediated neurodegeneration are not completely understood.

The isoforms of JNK react to the various external stimuli such as inflammatory, interleukins, tumor necrosis factor (TNF-α) and stress in different ways. Various studies have shown that the importance of JNK pathway playing a key role in the neurodegeneration in glaucoma and in induced animal models by ischemic injury, optic nerve crush ^[20, 34-36]^. In the photocoagulation method of IOP elevation, RGC loss was accompanied by JNK activation, suggesting the role of JNK in RGC apoptosis ^[37, 38]^. In an optic nerve crush model, the combined deletion of JNK2 and JNK3 and the conditional deletion of JUN inhibited the loss of RGC and showed the long-term protection ^[20]^ and the upstream blockade of the JNK signaling also enhanced the survival of RGC ^[34]^. In an ischemia/reperfusion model, administration of the JNK inhibitor SP600125 demonstrated the protective role in RGC as well as in non RGC cells in various inner retinal layers ^[35]^.

In our present study we focused on the involvement of JNK2 in ET-1 mediated RGC loss in mice. We used intravitreal administration of 1 nmole of ET-1 which was the dose that produced a significant decline in axonal transport following intravitreal administration in prior experiments in rats ^[39]^. To assess the increased phosphorylation of c-Jun we used the antibodies which can recognize the phosphorylation at Ser 63/73 residue, the specific site targeted by the kinase activity of JNK ^[40]^. Though the phosphorylation of c-Jun could occur through the actions of any of the three JNK isoforms, JNK2^-/-^ mice didn’t show the significant up regulation in the RGC layer, when compared to the wild type mice in both the 2 h and 24 h time point. Interestingly, we observed the significant upregulation of phospho c-Jun in GCL for 2 h and 24 h time point and in INL at 24 h time point. The relevance of changes in c-Jun phosphorylation in the INL is not clear. The comparison of the ET-1 treated eyes between the JNK2^-/-^ mice and wild type showed the significant upregulation of phospho-c-Jun in the wild type eyes in both RGC and non RGC layers (**Figure 1** and **Figure 2**).

To further determine whether the upregulation of phospho c-Jun could lead to the RGC death we assessed RGC survival by counting Brn3a-positive cells in retinal flat mounts. We observed that a significant (p=0.0082, n=6) an overall 26% loss of RGCs was found in the retina in wild type mice, 7 days post injection with ET-1, compared to that in vehicle injected wild type mice. JNK2^-/-^ mice showed no significant (p=0.36, n=5) loss of RGC after ET-1 administration, compared to JNK2^-/-^ mice treated with the vehicle (**Figure 3**). This data clearly shows the neuroprotective role of JNK2 in RGC with the administration of ET-1. It is possible that JNK2 plays an important role in signal transduction mechanisms following ET-1 mediated neurodegeneration, since phosphorylation of JNK is significantly attenuated in JNK2 knockout mice. Moreover, ET-1-mediated phosphorylation of c-Jun is greatly reduced in JNK2^-/-^ mice and there is no compensatory phosphorylation by JNK-1 and JNK-3 in the soma of RGCs.

In glaucoma and many neurodegenerative diseases, the axonal degeneration often leads to the apoptotic of the soma. Studies in genetic mouse model by over expression of Bcl2, showed the prevention of soma death but not the degeneration of axons ^[41,42]^. Most of the studies in rodent models with induced ocular hypertension have shown that axonal degeneration and dysfunction precedes RGC death, early in course of disease progression, which ultimately leads to the axonal transport obstruction, thereby leading to the RGC loss ^[43-49].^In C57BL/6 background mice, the combined null mutation of JNK 2 and JNK3 protected dopamine neurons, however was not able to protect against axon degeneration, following intrastriatal administration of 6-hydroxydopamine ^[50]^. On similar grounds, DBA 2J mouse model of ocular hypertension, the knockout of JNK2 and JNK3 protected the somas of RGCs, but not the axons from the degeneration ^[51]^. However, Libby’s group have shown that JNK2/JNK3 double knockout mice show significant protection from axonal degeneration following optic nerve crush in mice. Presently in our studies we also observed that the JNK2 deletion delayed the apoptotic RGC loss after the intravitreal injection administration of ET-1, but did not block axonal degeneration in mice.

In summary, ET-1 binding to its receptor could activate the JNK signaling mechanism thereby phosphorylating its downstream transcription factors. The extrinsic pathway of apoptosis which can cause RGC loss is mainly by up regulating the transcription factor, phospho c-JUN which is the immediate early gene response by the JNK. Our data suggest that JNK2 plays an important role in ET-1 mediated phosphorylation of c-Jun which can lead to the loss of RGCs, a key event in glaucoma pathology.

## Declarations

### Authors contributions

B.K, R.K designed research studies, analyzed data, performed experiments and wrote the manuscript. D.S assisted in conducting key experiments. V.K assisted in imaging and data analysis. All authors read and approved the final manuscript.

### Ethics approval

All experiments and procedures performed in this study were in agreement with the ARVO guideline for the use of animals in research and were approved by the Institutional Animal Care and Use Committee (IACUC) at the UNT Health Science Center, Fort Worth.

### Consent for publication

All authors have reviewed the manuscript and given their consent for publication.

### Availability of data and material

Not applicable.

